# Genetic diversity of *Staphylococcus aureus* wall teichoic acid glycosyltransferases affects immune recognition

**DOI:** 10.1101/2022.05.25.492469

**Authors:** Sara M. Tamminga, Simon L. Völpel, Kim Schipper, Thilo Stehle, Yvonne Pannekoek, Nina M. van Sorge

## Abstract

*Staphylococcus aureus* (*S. aureus*) is a leading cause of skin and soft tissue infections and (hospital-acquired) systemic infections. Wall teichoic acids (WTAs) are cell wall-anchored glycopolymers that are important for *S. aureus* nasal colonization, endocarditis, and antibiotic resistance. WTAs consist of a polymerized ribitol phosphate (RboP) chain that can be glycosylated with *N*-acetylglucosamine (GlcNAc) by three glycosyltransferases: TarS, TarM, and TarP. TarS and TarP modify WTA with β-linked GlcNAc on the C-4 (β1,4-GlcNAc) and the C-3 position (β1,3-GlcNAc) of the RboP subunit, respectively, whereas TarM modifies WTA with α-linked GlcNAc at the C-4 position (α1,4-GlcNAc). Importantly, these WTA glycosylation patterns impact immune recognition and clearance of *S. aureus*. Previous studies suggest that *tarS* is near-universally expressed within the *S. aureus* population, whereas a smaller proportion co-express either *tarM* or *tarP*. To gain more insight in the presence and genetic variation of *tarS, tarM*, and *tarP* in the S. aureus population, we analyzed a collection of 25,652 *S. aureus* genomes within the PubMLST database. Over 99% of isolates contained *tarS*. Co-expression of *tarS*/*tarM* or *tarS*/*tarP* occurred in 37% and 7% of isolates, respectively, and was associated to specific *S. aureus* clonal complexes. We also identified 26 isolates (0.1%) that contained all three glycosyltransferase genes. At sequence level, we identified *tar* alleles with amino acid substitutions in critical enzymatic residues or with premature stop codons. Several *tar* variants were expressed in a *S. aureus tar*-negative strain. Analysis using specific monoclonal antibodies and human langerin showed that WTA glycosylation was severely attenuated or absent. Overall, our data provide a broad overview of the genetic diversity of the three WTA glycosyltransferases in the *S. aureus* population and the functional consequences for immune recognition.

## Introduction

*Staphylococcus aureus* (*S. aureus*) is a common member within microbiota communities and colonizes approximately 30% of the human population asymptomatically^1^. However, *S. aureus* is also a prominent bacterial pathogen in hospital- and community-acquired infections. Infections often start locally, for example in the skin, but if local immune recognition and immune defense fail, bacteria can disseminate and cause systemic infections. Even with timely treatment and clinical management, such infections are associated with high overall disease burden and mortality.

A major component of the *S. aureus* cell wall is wall teichoic acid (WTA), which is important for nasal colonization, β-lactam resistance, and phage-mediated horizontal gene transfer^2^. WTA molecules are composed of 20-40 ribitol phosphate (RboP) subunits that are polymerized into a linear backbone, which is modified with D-alanine and *N*-acetylglucosamine (GlcNAc) moieties^3^. Currently, three different glycosyltransferases, *i*.*e*. TarS, TarM and TarP, are known to glycosylate WTA with GlcNAc^3-8^. GlcNAc is attached to the C-4 hydroxyl group of RboP in either the α- or β-configuration by TarM and TarS, respectively^3, 8^. Similar to TarS, TarP attaches GlcNAc in a β-configuration, but to the C-3 hydroxyl group of RboP^6^. All three enzymes have been characterized both functionally and structurally, providing insight on critical protein residues and features for catalysis^4-7^.

The specific WTA GlcNAc modifications greatly influence β-lactam- and phage resistance of *S. aureus*^2, 3^ as well as host-pathogen interaction^9-12^. For instance, the WTA glycan modifications represent dominant antigens in the *S. aureus*-reactive antibody pool in humans^2, 9, 12^. Most antibodies are directed against β-GlcNAc modified WTA^9^ and not every WTA glycoform may be similar in terms of immunogenicity or as target for antibody-mediated clearance^6, 9^. With regard to innate immunity, β-GlcNAc-WTA but not α-GlcNAc-WTA is detected by the innate receptor langerin, which is expressed on skin epidermal Langerhans cells^10, 11^. These findings indicate that different WTA glycosylation profiles, which depend on the presence and activity of the three known glycosyltransferases, impact *S. aureus* immune recognition and clearance.

Previous studies suggest that *S. aureus* near-universally expresses *tarS*, whereas a smaller proportion co-expresses either *tarM* or *tarP*^1, 6, 13^. Indeed, *tarS* is part of the *S. aureus* core genome and clusters within well-studied WTA biosynthesis genes^3^. In contrast, *tarM* is located elsewhere in the genome and is suggested to be an ancient genetic trait of *S. aureus*, based on the observation that strains belonging to the very early branching *S. aureus* bear both *tarS* and *tarM* in their genome^2, 13^. It is hypothesized that *tarM* was lost during *S. aureus* evolution, resulting in its absence in strains belonging to clonal complex (CC) 5 and CC398^13^. The most recently identified enzyme, *tarP*, is encoded on different prophages and its presence seems restricted to isolates belonging to CC5 and CC398^6^. However, an overview on the presence, co-expression, and genetic variation of *tarS, tarM* and *tarP* within the *S. aureus* population is currently lacking.

In this study, we dissected the presence and genetic variation of *tarS, tarM*, and *tarP* in a collection of 25,652 *S. aureus* genomes that are deposited in the open-access PubMLST database (https://pubmlst.org/organisms/staphylococcus-aureus). We also analyzed whether specific combinations or *tar* allelic variants were associated with specific CC of *S. aureus*. Finally, we performed *in silico* analyses followed by cloning and plasmid-expression of several *tar* variants in a *S. aureus tar*-deficient strain to observe the functional effect of *tar* sequence variation on WTA glycosylation using WTA-specific Fab fragments and human langerin. Overall, our data provide more insight into the genetic diversity of the three WTA GlcNAc-transferases and demonstrates how this natural variation can impact *S. aureus* immune recognition by both the innate and adaptive immune system. As such, our study provides a blueprint to dissect and functionally analyze *S. aureus* genes at a population-wide level.

## Material and methods

### *S. aureus* PubMLST database analysis

We analyzed the *S*.*aureus* genomes deposited in the PubMLST Bacterial Isolate Genome Sequence Database (BIGSdb) to determine the presence and genetic diversity of *tar*-glycosyltransferases *tarS, tarM*, and *tarP* (identified as SAUR2940, SAUR2942, and SAUR2941, respectively in the PubMLST *S. aureus* database). Of the 26,605 genomes in the database (extracted on 08-02-2022), several genomes were excluded from further analysis based on the following criteria. Fifty-seven isolates were excluded since they were suspected to be contaminated or not *S. aureus*, based on the comments section of the database. Furthermore, 691 isolates were excluded since they contained more than 300 contigs and/or had a N50 contig length of <20,000 bp. Strains were also excluded when a *tar*-gene was not present within a single contig (n=205). After exclusion of these strains, 25,652 *S. aureus* isolates remained for analysis. Allele numbers and nucleotide sequences of the three glycosyltransferases from these isolates were downloaded from the PubMLST *S. aureus* BIGSdb. To verify that all isolates that contained a *tar-*gene also had an assigned allele in the database, an additional BLAST analysis was performed using allele 1 for *tarS, tarM*, and *tarP* as reference genes. Isolates that contained a *tar-*gene sequence but lacked an assigned allele were scanned with 70% stringency and novel alleles were assigned and added to the database. In the case of a premature stop codon in *tar* sequence, the gene was marked as ‘truncated’, and no allele was assigned. Information on the isolates’ sequence type (ST) and CC was also obtained from the PubMLST *S. aureus* BIGSdb. For some isolates the CC in the database was not defined, therefore we manually added CC7, CC12, CC25, CC59, CC88, CC130, CC133, CC398, and CC425 based on ST information^14, 15^.

### Alignments and protein sequence analysis

The nucleotide sequences of all allelic variants of *tarS, tarM*, and *tarP* were translated to amino acid sequences and aligned using MUSCLE in MEGA version 11^16^. We manually analyzed critical residues of the enzymes (based on previous studies)^4-7^ for amino acid substitutions in or ± 1 amino acid next to the critical residues of Tar-enzymes. The effect of the amino acid substitutions on interactions with the acceptor substrate (poly-RboP) or the donor substrate (UDP-GlcNAc) was predicted and visualized using the PyMOL Molecular Graphics System, Version 2.5.1 Schrödinger, LLC. During visualization, we continuously chose the rotamer orientation for the mutated residue that had the least strains and clashes with other residues in the structure.

### Bacterial strains and culture conditions

All plasmids and *S. aureus* strains used for wet-lab experiments in this study are listed in Table S1. Bacteria were grown overnight in 5 mL tryptic soy broth (TSB, Oxoid) at 37 °C with agitation. For *S. aureus* strains that were complemented with plasmid, TSB was supplemented with 10 μg/mL chloramphenicol (Sigma). Overnight cultures were subcultured the next day in fresh TSB and grown to mid-exponential growth phase, corresponding to an optical density of 0.6 to 0.7 at 600 nm (OD_600_).

### Generation of complemented RN4220 Δ*tarM*S strains with *tar* variants

Shuttle vector RB474^17^ containing full-length copies of *tarS, tarM*, or *tarP* as inserts (Table S2) were used to recreate naturally-occurring amino acid substitutions and premature stopcodons in *tar*-glycosyltransferases ^3, 6, 8^. Nucleotide substitutions were generated using either QuikChange Site-Directed Mutagenesis Kit (Agilent) or the required nucleotide sequence was ordered as gBlock (IDT). Used primers and gBlocks are listed in Table S3. Plasmids containing sequences of *tarS, tarM*, or *tarP* were amplified in *E. coli* DC10b^18^ or *E. coli* XL-1 and transformed into electrocompetent *S. aureus* RN4220 Δ*tarMS*^*19*^ through electroporation with a Bio-Rad Gene Pulser II (2.0 kV, 600 Ω, 10 μF). After recovery, bacteria were plated on tryptic soy agar (TSA) plates supplemented with 10 μg/mL chloramphenicol to select plasmid-complemented colonies. The presence of *tarS, tarM*, or *tarP* was confirmed by polymerase chain reaction (PCR) analysis using *tar* gene-specific primers and nucleotide sequences of the constructs were confirmed by Sanger sequencing.

### Analysis of WTA glycosylation using monoclonal Fab fragments and human langerin-FITC

The enzymatic activity of the *tar-*variants was assessed by analyzing WTA glycosylation using WTA-specific Fab fragments^11^. Bacteria were grown to mid-exponential growth phase, collected by centrifugation (4,000 rpm, 8 min, 4°C) and resuspended at an OD_600_ of 0.4 (∼1× 10^8^ colony forming units (CFU)/mL) in phosphate-buffered saline (PBS; pH 7) with 0.1% bovine serum albumin (BSA; Sigma). Bacteria (1.25 × 10^6^ CFU) were incubated with 3.3 μg/mL monoclonal Fab fragments (pre-diluted in PBS 0.1% BSA) specific to β-GlcNAc (clone 4497) or α-1,4-GlcNAc (clone 4461) WTA, respectively^11^. After washing, bacteria were incubated with a Goat F(ab’)_2_ anti-human Kappa-Alexa Fluor 647 (pre-diluted in PBS 0.1%BSA; 2.5 μg/mL, Southern Biotech #2062-31) to detect bound Fab fragments. Bacteria were washed and fixed in PBS with 1% paraformaldehyde, and analyzed by flow cytometry on a BD FACSCanto II Flow Cytometer (BD Bioscience). Per sample, 10,000 gated events were collected and fluorescence was expressed as geometric mean. Additionally, bacterial binding to recombinant FITC-labeled human langerin (kindly provided by Prof. C. Rademacher, University of Vienna, Vienna, Austria) was assessed as previously described^10^. Briefly, bacteria were grown to mid-exponential phase, collected, and resuspended at an OD_600_ of 0.4 in TSM buffer (2.4g/L Tris (Sigma-Aldrich), 8.77 g/L NaCl (Merck), 294 mg/L CaCl_2_(H_2_O)_2_ (Merck), 294 mg/L MgCl_2_(H_2_O)_6_ (Merck), containing 0.1% BSA (Sigma), pH = 7). Next, bacteria were incubated shaking for 30 min at 37°C with 20 μg/mL FITC-labeled human langerin-extracellular domain (ECD) constructs (referred to as langerin-FITC). Finally, bacteria were washed once with TSM 0.1% BSA, fixed in 1% paraformaldehyde in PBS and analyzed by flow cytometry as described above.

### Statistical analysis

Flow cytometry data were analyzed using FlowJo 10 (FlowJo, LLC). All data were analyzed using GraphPad Prism 9.1.0 (GraphPad Software) with a one-way ANOVA followed by a Dunnett’s multiple comparison test. *p*-Values are depicted in the figures, and *p* < 0.05 was considered significant.

## Results

### *TarS* is present in >99% of all *S. aureus* isolates

Analysis of 25,652 *S. aureus* genomes demonstrated that over 99% of isolates contained *tarS*, with 36.7% and 6.6% of isolates co-expressing *tarM* (n= 9,404) or *tarP* (n=1,702), respectively (Table 1). We also found 26 isolates (0.10%) that contained all three glycosyltransferase genes (Table 1). Overall, to 186 isolates no *tarS* alleles could be assigned in the following combinations; to 67 isolates (0.26%) no *tar* alleles could be assigned, to 58 isolates (0.23%) only a *tarP* allele could be assigned, to 56 isolates (0.22%) only a *tarM* allele could be assigned and to 5 isolates (0.019%) both *tarP* and *tarM* alleles could be assigned (Table 1). Further analysis of these strains lacking a *tarS* allele identified an incomplete open reading frame (ORF) for *tarS* in 165 out-of-186 isolates (88.7%) due to the presence of a premature stop codon. We also found premature stop codons in sequences of *tarM* (n=106) and *tarP* (n=2). Isolates with premature stop codons in specific *tar* genes are indicated as ‘truncated’ in Table 1.

**Table 1.** Presence of *tar*S, *tarM* and *tarP* in 25,652 *S. aureus* isolates

| *tar* genotype | Isolates (n) | Percentage of total |
| --- | --- | --- |
| *tarS* | 14,334 | 55.9 |
| *tarS* + truncated *tarM* | 98 |  |
| *tarS* + truncated *tarP* | 2 |  |
| *tarS* only | 14,234 |  |
| *tarS* + *tarM* | 9,404 | 36.7 |
| *tarS* + *tarP* | 1,702 | 6.6 |
| *tarS* + *tarP* + truncated *tarM* | 4 |  |
| *tarS* + *tarP* | 1,698 |  |
| No *tar* alleles assigned | 67 | 0.26 |
| truncated *tarS* | 57 |  |
| truncated *tarM* | 4 |  |
| No *tar* genes | 6 |  |
| *tarP* | 58 | 0.23 |
| *tarP* + truncated *tarS* | 58 |  |
| *tarM* | 56 | 0.22 |
| *tarM* + truncated *tarS* | 45 |  |
| *tarM* only | 11 |  |
| *tarS* + *tarM* + *tarP* | 26 | 0.10 |
| *tarM* + *tarP* | 5 | 0.02 |
| *tarP* + *tarM* + truncated *tarS* | 5 |  |
| **Total** | **25,652** | **100** |

### Presence of *tar*-glycosyltransferases is linked to *S. aureus* Clonal Complexes

Next, we analyzed whether *tar* genotype was associated with specific CC of *S. aureus*. The most prevalent CCs in the database were CC8 (26%), CC5 (23%), and CC22 (20%) (Figure 1A), which are all CCs associated with human disease including MRSA strains^15, 20-22^. Besides being a successful human pathogenic lineage, CC5 is also often found in poultry infections^14, 23^. Other livestock- and animal-associated CCs that were present in the database include CC97, CC130, CC133, CC398, and CC425^14, 15, 21, 23^. Of the isolates that only contained a complete *tarS* gene, 36% belonged to CC22 and 32% to CC5 (Figure 1B). Overall, the *tarS/tarM* combination was predominantly found in isolates belonging to CC8 (Figure 1C), even when *tarP* (Figure 1G and 1H) or a truncated version of the *tarS* gene was present (Figure 1I). In total, *tarM* was found in 7 different CCs (Table S4). For isolates that contain *tarS* and *tarP*, nearly all belonged to CC5 (69%) or CC398 (23%) as previously described^6^ (Figure 1D). Yet, *tarP* was also identified in at least 11 additional CCs (Table S4). Interestingly, of the 58 isolates that only contained complete *tarP* gene, 52% belonged to CC45 (Figure 1F), whereas this CC represented only 0.4% of the *tarS*/*tarP*-containing isolates.

**Figure 1.**
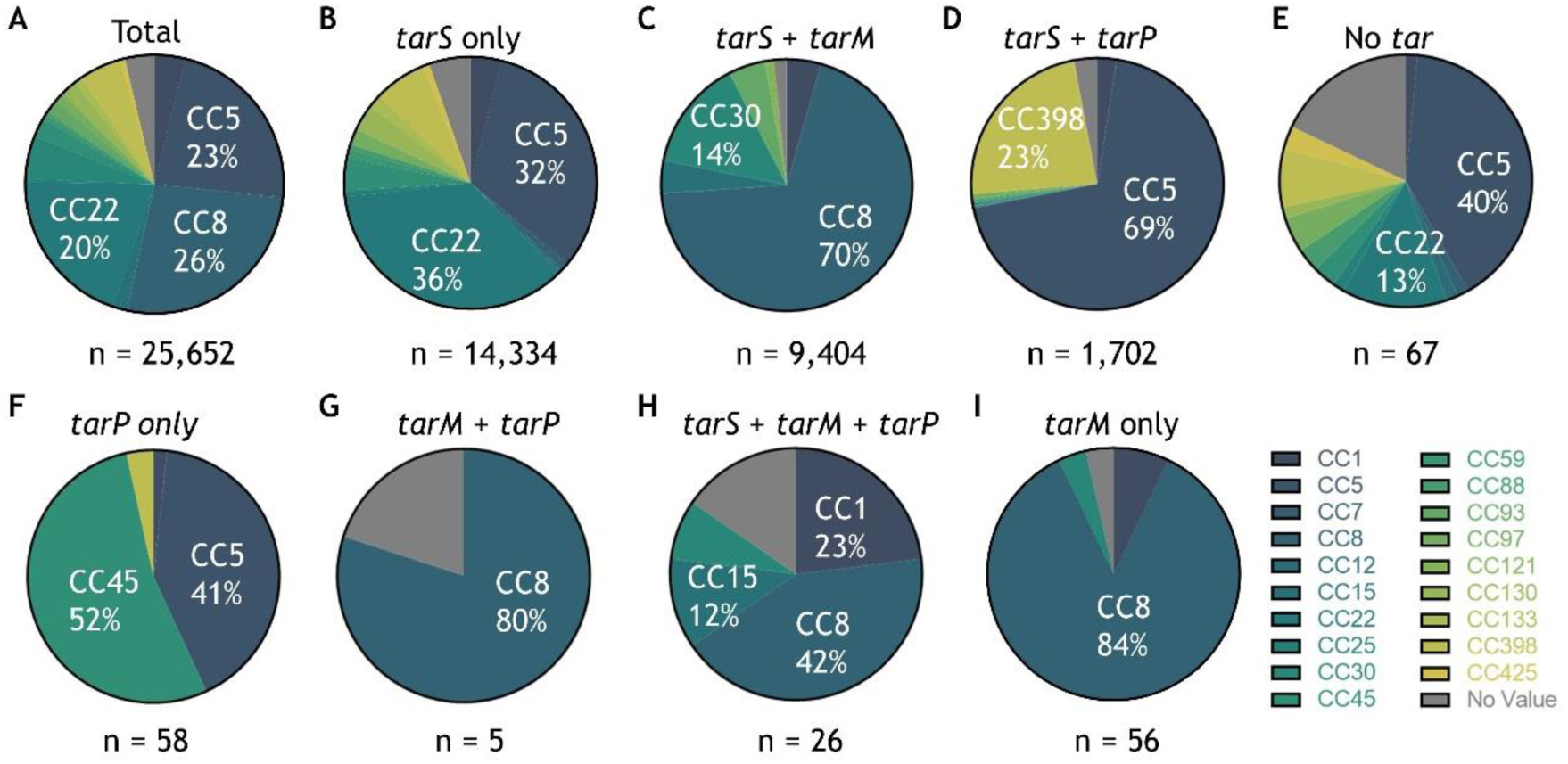
*tar* genotype is linked to *S. aureus* clonal complexes. (A) Distribution of CCs for isolates within the PubMLST database on the extraction date. Only CCs that comprise of >10% of isolates are indicated. Color coding of CCs is shown in legend on the right. (B-I) *S. aureus* CC distribution is shown for each *tar* genotype indicated as in (A).

### *Tar* alleles and CC distribution

Based on nucleotide sequence, 536 alleles (coding for 408 unique ORFs) were identified for *tarS* within the PubMLST *S. aureus* database, and 8 alleles (coding for 6 unique ORFs) comprised 77% of all *tarS* nucleotide sequences (Figure 2A). The number of alleles was much smaller for *tarM* with 180 nucleotide sequences coding for 143 unique ORFs, and *tarP* with only 28 alleles coding for 19 unique ORFs. For these genes, allele 1 was most frequently found, representing 71% of all *tarM* (Figure 2B) and 71% of all *tarP*-sequences (Figure 2C). All CC, with the exception of *tarS* for CC1 and 8, were characterized by the presence of a single dominant allele of *tar-*glycosyltransferases (Figure S1).

**Figure 2.**
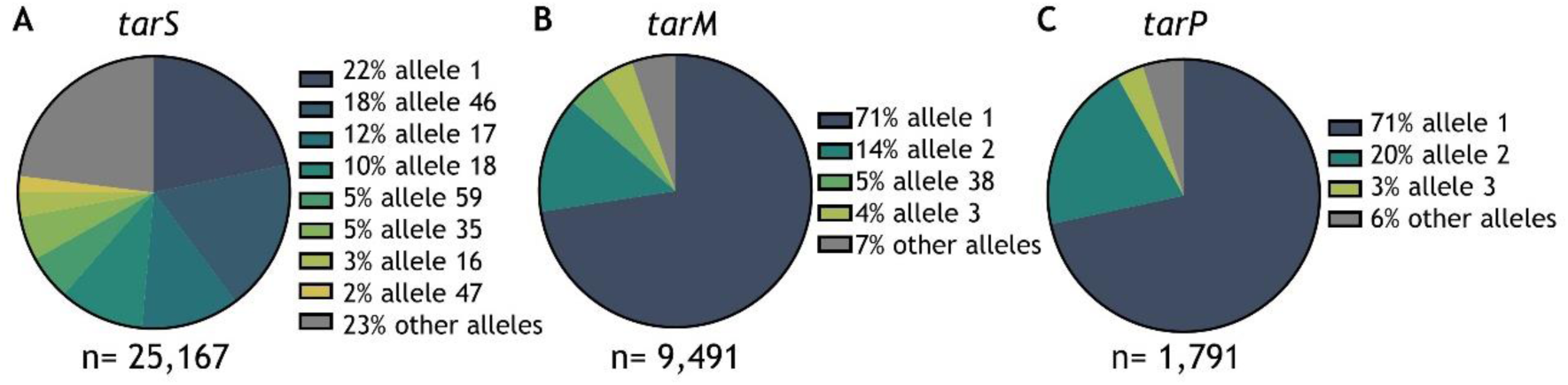
Allelic variants of *tarS, tarM*, and *tarP* in the PubMLST *S. aureus* collection. (A) Representation of the dominant *tarS* alleles within the *S. aureus* isolates of the PubMLST database. Only alleles that cover >1% of all sequences are displayed individually, remaining minor alleles are grouped (grey). Total amount of alleles (based on nucleotide sequence) identified for *tarS* was 536. (B) Same as in (A) but for *tarM*, total amount of alleles was n=180. (C) Same as in (A) but for *tarP* total amount of alleles was n=28.

### Amino acid substitutions in critical residues of Tar-glycosyltransferases

We identified several *tar* alleles with a naturally-occurring amino acid substitution in a critical residue of the enzyme, based on information from the published crystal structures ^5-7^. Overall, these alleles were identified in 32 isolates (0.12%) of the analyzed *S. aureus* genomes. The amino acid changes in these variants and the function assigned to these residues are listed in Table 2. Figure 3 visualizes the amino acid substitutions and their predicted effect on molecular interactions in PyMOL. D91 is part of the DxD motif in TarS, which is directly involved in the interaction with UDP-GlcNAc^5^ (depicted as black dashed line; Figure 3A). Substitution of an acidic aspartate for a basic histidine, which also has a bulkier side chain, is predicted to affect UDP-GlcNAc binding (Figure 3A). TarS E177 makes contact with the C4-OH and C6-OH of the UDP-GlcNAc moiety^5^ (black dashed line; Figure 3B) and supports orientation of D178 (base catalyst). Consequently, substitution of E177 to K177 is likely to conflict with P71 (slate; Figure 3B) in addition to a change from a positive-to a negatively-charged residue. Moreover, the distance of the terminal primary amine group to UDP-GlcNAc C4-OH and C6-OH becomes 4.5 Å (compared to 2.7 Å and 3.0 Å of the E177 carboxy group to UDP-GlcNAc C4-OH and C6-OH, respectively), which may hamper sufficient hydrogen bonding (Figure 3B). A third mutation in *tarS* results in substitution of S212 to R212 (Figure 3C). TarS S212 coordinates the β-phosphate of the donor substrate UDP-GlcNAc and is positioned in close proximity to both the donor and acceptor substrate binding sites^5^. During the amino acid substitution (S212R) visualization in PyMOL, we could place several rotamers that did not conflict with TarS structure (Figure 3C). However, all these rotamers point to the spaces between the two sulfates (yellow; Figure S2) which might indicate the phosphate binding sites of poly-RboP and therefore may interfere with binding of the acceptor substrate. Furthermore, the coordination of the β-phosphate might be lost as arginine is too long to facilitate this (Figure 3C). For TarM, the backbone nitrogen atom of G17 coordinates the α-phosphate of UDP-GlcNAc^7^. Consequently, substitution of G17 to D17 is predicted to affect the interaction of TarM with UDP-GlcNAc as there appears to be little space for the aspartate side chain (Figure 3D). In TarP, no amino acid substitutions of critical residues were identified, but we did identify an amino acid substitution (T10P) next to a critical residue (F11). F11 stabilizes the uracil moiety of UDP with its aromatic sidechain and interacts with the backbone carbonyl of the ribose moiety^6^, and mutation of the neighboring amino acid may affect this interaction (Figure 3E). Moreover, T10P forms a proline-proline peptide bond with P9, and this diproline segment is even more restricted in its conformational arrangement compared to a single proline (Figure 3E).

**Table 2.** Naturally-occurring amino acid substitutions in *tarS, tarM* and *tarP* with function of original residue

| ***tar* gene** | **Substitution** | **Isolates (n)** | **Function residue** |
| --- | --- | --- | --- |
| *tarS* | D91H | 14 | Interaction with C3-OH of GlcNAc<sup>5</sup> |
|  | E177K | 6 | Interaction with C4-OH and C6-OH of GlcNAc<sup>5</sup> |
| | S212R | 3 | Coordinates β-phosphate of UDP-GlcNAc<sup>5</sup> |
| *tarM* | G17D | 2 | Interaction with α-phosphate of the UDP-GlcNAc<sup>7</sup> |
| *tarP* | T10P | 7 | F11 is important for GlcNAc interaction<sup>6</sup> |

**Figure 3.**
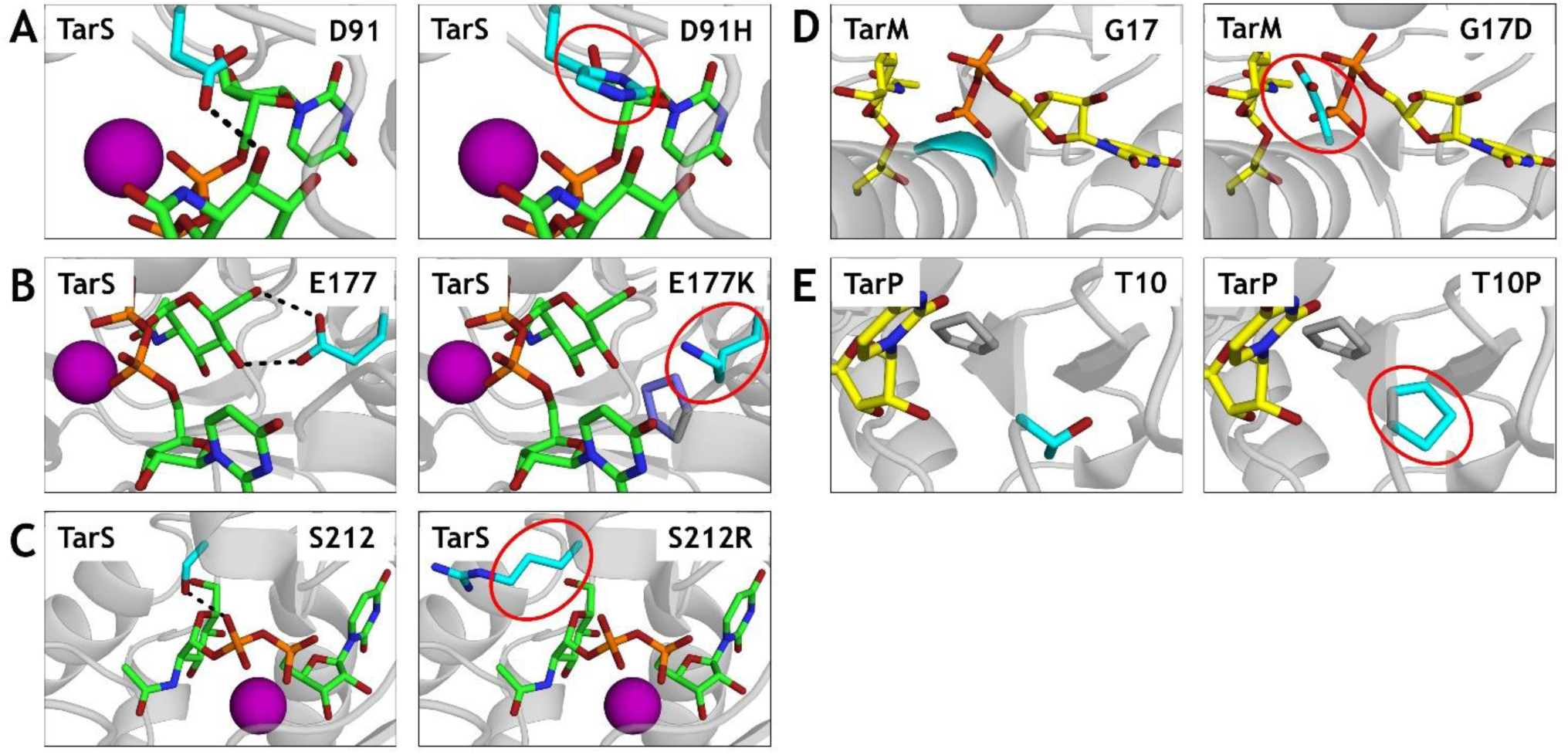
PyMOL visualizations of the effect of naturally-occurring amino acid substitutions on molecular interactions within Tar-enzymes. (A) TarS is shown in grey cartoon presentation (PDB code 5TZE). The donor substrate UDP-GlcNAc (green) and the residues of interest: unaltered (left) and the substitution (right, circled red), both in cyan, are all displayed in stick form. Black dashed line indicates interaction with UDP-GlcNAc. The Mn^2+^ ion coordinating the pyrophosphate of UDP is presented as sphere in magenta. (B) In the E177K mutation (right) the conflicting residue P71 is displayed in stick (slate). (C) TarS, S212 and S212R are displayed as in (A). (D) TarM is shown in cartoon presentation in grey (PDB code 4X7R). UDP and α-glyceryl-GlcNAc are displayed in stick (yellow). Rest of residues are colored as in (A). (E) TarP, T10 and T10P are shown as in (D) with P9 (grey) as stick model (PDB code 6H4M)

### Premature stopcodons found in *tar*-glycosyltransferases genes

Next, we furthermore wanted to predict the impact of the naturally-occurring premature stop codons (indicated as truncated in Table1). Therefore we visualized the location of the stop codons in 2D-scaled models of *tar*-glycosyltransferases based on previously published enzyme structures^4-7^ (Figure 4). *TarS* (1,719 nt) contains two domains that are important for GlcNAc interaction (nt 28-393 and 529-636; yellow) and a poly-RboP interaction domain (nt 742-888; red). Furthermore, TarS contains two residues that are associated with trimerization (nt 1561-1563 and 1594-1596; blue). Certain premature stop codon positions were found in 5 or more (up to 29) different isolates (Figure 4A). A similar schematic overview is shown for *tarM* (Figure 4B) and *tarP* (Figure 4C). Premature stop codons in *tarM* (1,506 nt) were identified in 106 isolates (Figure 4B). For *tarP* (984 nt), only 2 isolates contained a premature stop codon (Figure 4C). Overall, premature stopcodons were found across the entire length of *tarS* and *tarM* genes and may affect enzymatic functionality depending on their position within the enzyme.

**Figure 4.**
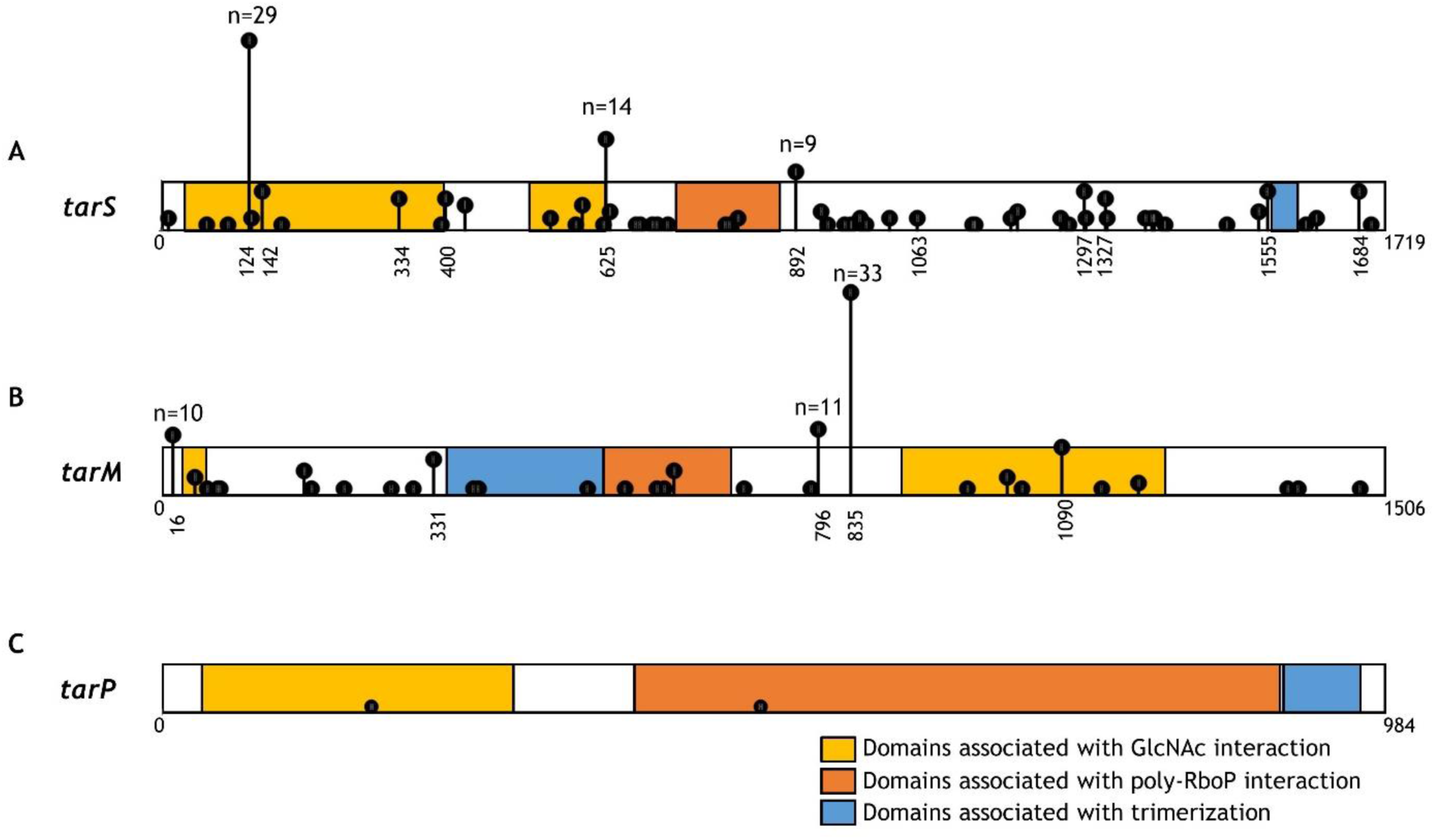
*Tar-*glycosyltransferase genes depicted as scaled-2D model with nucleotide position of premature stop codons. (A) Scaled 2D representation of *tarS* (nt 1,719) containing two domains important for GlcNAc interaction (nt 28-393 and 529-636; yellow), a poly-RboP interaction domain (nt 742-888; orange) and two residues associated with trimerization (nt 1561-1563 and 1594-1596; blue). Premature stop codons were identified in 165 isolates and are indicated by the vertical black lines that show position and frequency. For premature stop codons that were present in >4 isolates, nt position is shown and for >8 the exact number of isolates is indicated. (B) *tarM* (1,506 nt) contains two residues at the start (nt 49-54; yellow) and domain (nt 910-1233; yellow) that are important in GlcNAc interaction. Domain 349-540 (blue) contains the HUB domain (formerly known as DUF1975) associated with TarM trimerization. Directly adjacent is the poly-RboP interaction domain (nt 543-699; orange). Premature stop codons (n=106 isolates) are indicated as in (A). (C) *tarP* (984 nt) contains a domain associated with GlcNAc interaction (nt 31-285; yellow) followed by a poly-RboP interaction domain (385-789; orange). Domain 916-978 (blue) is associated with trimerization of the enzyme. Premature stop codons (n=2 isolates) are indicated as in (A).

### Amino acid substitutions and premature stop codons hamper *S. aureus* immune recognition by antibodies and langerin

The functional effect of naturally-occurring amino acid substitutions and premature stop codons in *tar*-glycosyltransferases on WTA-GlcNAc decoration, was assessed by expressing these *tarS, tarM* and *tarP* variants in a *S. aureus* mutant lacking WTA glycosylation (Δ*tarMS*). As a read-out of WTA glycosylation, we used specific Fab fragments against β-GlcNAc-WTA (clone 4497) and α-1,4-GlcNAc-WTA (clone 4461)^11^. Furthermore, we recently identified that β-GlcNAc-WTA is specifically detected by the human innate receptor langerin^10^. Therefore, we also determined bacterial binding of the *tar* variants by langerin-FITC. For TarS we tested the effect of a premature stop codon on nucleotide positions 124, 625, 892 and 1063 (Figure 4A). Overall, β-GlcNAcylation of WTA is significantly (*p*<0.0001) decreased in all four premature stopcodon variants of TarS compared to wild-type (WT) TarS but to varying extend. Premature stop codons on nucleotide positions 124 and 625 abrogated interaction with β-GlcNAc specific Fab fragments and langerin (Figure 5A,B). Closer to the C-terminus of TarS, stopcodons on nucleotide positions 892 and 1063 were also severly hampered but still showed some residual activity, as β-GlcNAcylated WTA was still detectable with β-GlcNAc specific Fab fragments (Figure 5A. This level of glycosylation was insufficient for binding to langerin (Figure 5B). Next, we analyzed the effect of amino acid substitutions D91H, E177K and S212R in TarS and control mutations that were previously reported to attenuate TarS enzymatic activity *in vitro*^5^: D91A, E177A and S212A. Amino acid substitutions D91H and E177K in TarS completely abolished decoration of WTA with β-GlcNAc, similar to their controls D91A and E177A, as demonstrated by completely abrogated Fab binding (Figure 5C, *p*<0.0001) and langerin binding (Figure 5D, *p*<0.0001) compared to TarS WT (Figure 5D). Of note, amino acid substitution S212R also significantly (*p*<0.0001) reduced WTA β-GlcNAcylation and thereby immune recognition (Figure 5CD). Interestingly, control substitution S212A showed normal functionality, similar to WT TarS. TarP substitution T10P significantly (*p*<0.01) reduced WTA β-GlcNAcylation as well as langerin binding (*p*<0.01) compared to WT TarP (Figure 5EF). Lastly, WTA α-GlcNAcylation by TarM was completely abolished by the G17D amino acid substitution (*p*<0.0001) compared to WT TarM (Figure 5G). In conclusion, we showed that naturally-occurring premature stopcodons and amino acid substitutions can strongly affect WTA GlcNAcylation thereby affecting immune recognition by innate and the adaptive immune components.

**Figure 5.**
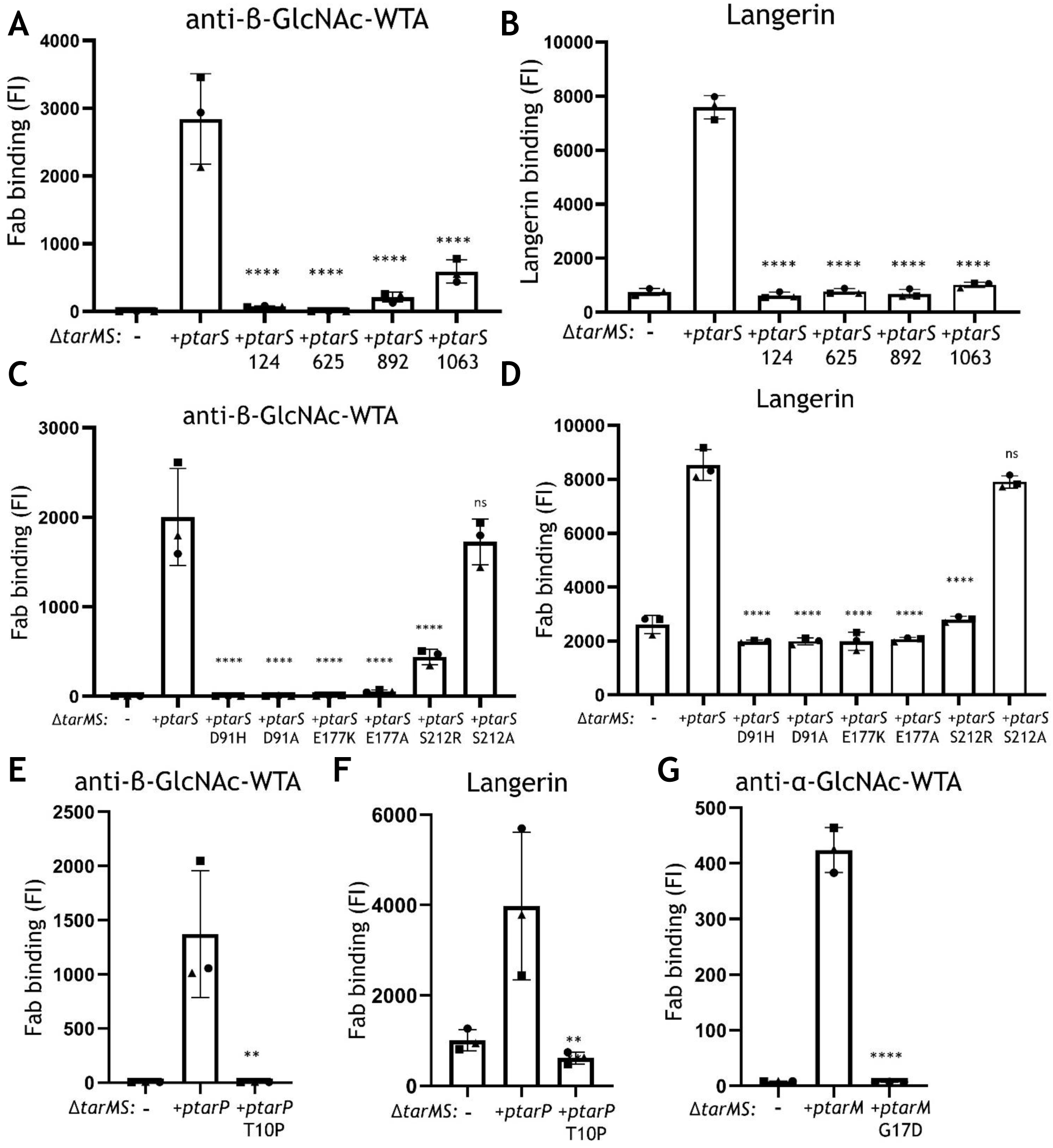
Impact of amino acid substitutions and premature stopcodons in TarS, TarM and TarP on *S. aureus* immune recognition. Binding of (A, C, E) monoclonal Fab fragments specific to β-GlcNAc-WTA (4497) and (B, D, F) human recombinant langerin-FITC to *S. aureus* RN4220 Δ*tarMS* complemented with plasmid-expressed WT *tarS* or premature stop codon *tarS* (A, B), *S. aureus* RN4220 Δ*tarMS* complemented with plasmid-expressed WT *tarS* or amino acid substitutions *tarS* variants (C, D) or *S. aureus* RN4220 Δ*tarMS* complemented with plasmid-expressed WT *tarP* or amino acid substitution *tarP* variant (E, F). (G) Binding of Fab fragments specific to α-GlcNAc-WTA to *S. aureus* Δ*tarMS* complemented with plasmid-expressed WT *tarM* or amino acid substitution *tarM* variant. Data are depicted as geometric mean fluorescence intensity (FI) of three individually displayed biological replicates + standard deviation (SD) and were compared to plasmid expressed WT enzyme \*\**p* < 0.01, \*\*\*\**p* < 0.0001.

## Discussion

GlcNAc decoration of *S. aureus* WTA by glycosyltransferases TarS, TarM and TarP is important for human nasal colonization, β-lactam resistance, phage-mediated horizontal gene transfer and immune recognition. Functional and structural analysis of *S. aureus* Tar-enzymes has been performed using a select number of strains and may not be representative for the entire *S. aureus* population. By analyzing 25,652 *S. aureus* genomes deposited in the PubMLST database, we confirmed that virtually all *S. aureus* isolates express *tarS* and that 37% and 7% of isolates co-express *tarM* and *tarP*, respectively. Co-expression of *tarS/tarM* or *tarS/tarP* correlated to specific *S. aureus* CCs. Moreover, we found a small number of *tar* alleles with natural amino acid substitutions in critical residues of the enzymes or with premature stop codons. By expressing these genes in a *tarMS*-deficient strain, we demonstrated that these genetic variants are severely attenuated in their enzymatic activity *in vivo*, thereby hampering immune recognition of *S. aureus* by innate and adaptive immune molecules.

Previous work on the presence of *tar*-glycosyltransferases only analyzed strain collections containing up to ∼100 isolates using PCR^2, 6, 13, 24, 25^. In this study, we analyzed 25,652 *S. aureus* genomes to obtain a more comprehensive overview of the distribution and genomic variability of the *tar*-glycosyltransferases across the *S. aureus* population. The PubMLST database contained isolates from 19 different CCs of human as well as animal origin, although there is clear skewing towards human clinical isolates. The lack of meta-data for many of the deposited genomes does not allow for a complete determination of their origin. Overall, the broad number of species and diversity of CCs renders PubMLST an important tool for *S. aureus* research on the presence and genetic variation of specific genes at the population level.

The presence of *tarP*, which is encoded on three different prophages^22^, was previously reported to be restricted to CC5 and CC398^6^. However, we also identified *tarP* in a small number of isolates from CC1, CC7, CC12, CC45, CC59, CC88, CC97, CC425, and even in *tarM-*associated CC8. This may suggest that *tarP* is present on more, at present unknown, phages, or that the host range of *tarP*-containing phages is broader than currently known.

Overall, our analysis showed that the *tar*-enzymes are highly conserved within the *S. aureus* population. Yet, amino acid substitutions in critical residues of the enzymes as well as premature stop codons, resulting in proteins without or with impaired enzymatic functionality, do occur. Consequently, the presence of one or more *tar* genes (identified by PCR or genome sequencing) does not provide complete information on WTA glycosylation in a particular isolate. In addition to gene presence and sequence, WTA glycosylation is also affected by environmental conditions^24^. Indeed, glycosylation by TarM and TarP is dominant over TarS during *in vitro* culture^6, 13^. For TarM this might be explained by a higher inherent enzymatic activity^5^, whereas for TarP this may be due to higher affinity for RboP compared to TarS^6^. Furthermore, *tar* genes may be transcriptionally regulated; *tarM* expression is increased during oxidative stress, most likely due to activation of the two-component GraRS regulon^26^. In contrast, *S. aureus* shifts towards TarS glycosylation at the expense of TarM/TarP WTA glycosylation during *in vivo* murine infection models and high salt conditions^24^. Overall, strain-specific WTA glycosylation will depend on specific gene sequence in combination with environmental-dependent gene expression and can only be assessed by direct staining methods such as specific Fab fragments.

We investigated the effect of specific genetic mutations on the enzymatic functionality by expressing these variants in a *tarMS*-deficient *S. aureus* strain. This analysis confirmed reduced or abolished activity for several specific amino acid substitutions and premature stop codons in TarS, TarM, and TarP. In addition to these naturally-occurring amino acid substitutions, we included amino acid substitutions in TarS (D91A, E177A and S212A) that were previously reported to attenuate TarS enzymatic activity *in vitro*^5^. However, in our FACS experiments using live *S. aureus* bacteria, we observed no significant differences in levels of β-glycosylated WTA with the S212A mutation compared to WT TarS, indicating that TarS enzymatic activity was similar to WT TarS. This may suggest that results for TarS enzymatic activity obtained *in vitro*, in which solely enzyme and substrate are present, are not always predictive for WTA β-glycosylation in live bacteria. However, the reason for this discrepancy remains to be elucidated. In addition, it should be noted that we only used the *S. aureus* RN4220 strain, which naturally contains *tarS* and *tarM*^*2*^, and effects may not be identical in different *S. aureus* strains.

In conclusion, *tar* glycosyltransferases are highly conserved and very abundant. Especially *tarS* was found to be present in >99% of *S. aureus* strains. We show that there are few exceptions in which *tar* genes seem present but contain amino acid substitutions or premature stop codons, for these isolates we show that *tar* genotype is not necessarily equal to *tar* phenotype and that thereby immune recognition is hampered. Studying the genetic presence and diversity of *tarS, tarM* and *tarP* provides more insight into *S. aureus* WTA glycosylation, which can help in development of anti-*S*.*aureus* preventive or therapeutic interventions such as monoclonal antibodies, phage therapy, and vaccines.

## Statements

### Data availability

Upon request, the data supporting these findings are available from the corresponding authors.

## Acknowledgments

The authors thank Dr. Carla J.C. de Haas and Dr. A. Robin Temming for the production of the monoclonal Fab fragments.

## Author contributions

Lab experiments: S.M.T, K.S. Data curation: S.M.T., Y.P. Software: S.L.V. Conceptualization: N.M.v.S., Y.P., T.S. Funding acquisition: N.M.v.S., T.S. Supervision: N.M.v.S., Y.P. Visualization: S.M.T, S.L.V. Writing – original draft: S.M.T. Writing – review & editing: N.M.v.S., Y.P., S.L.V.

## Supplemental Figures

**Supplementary Table 1.**
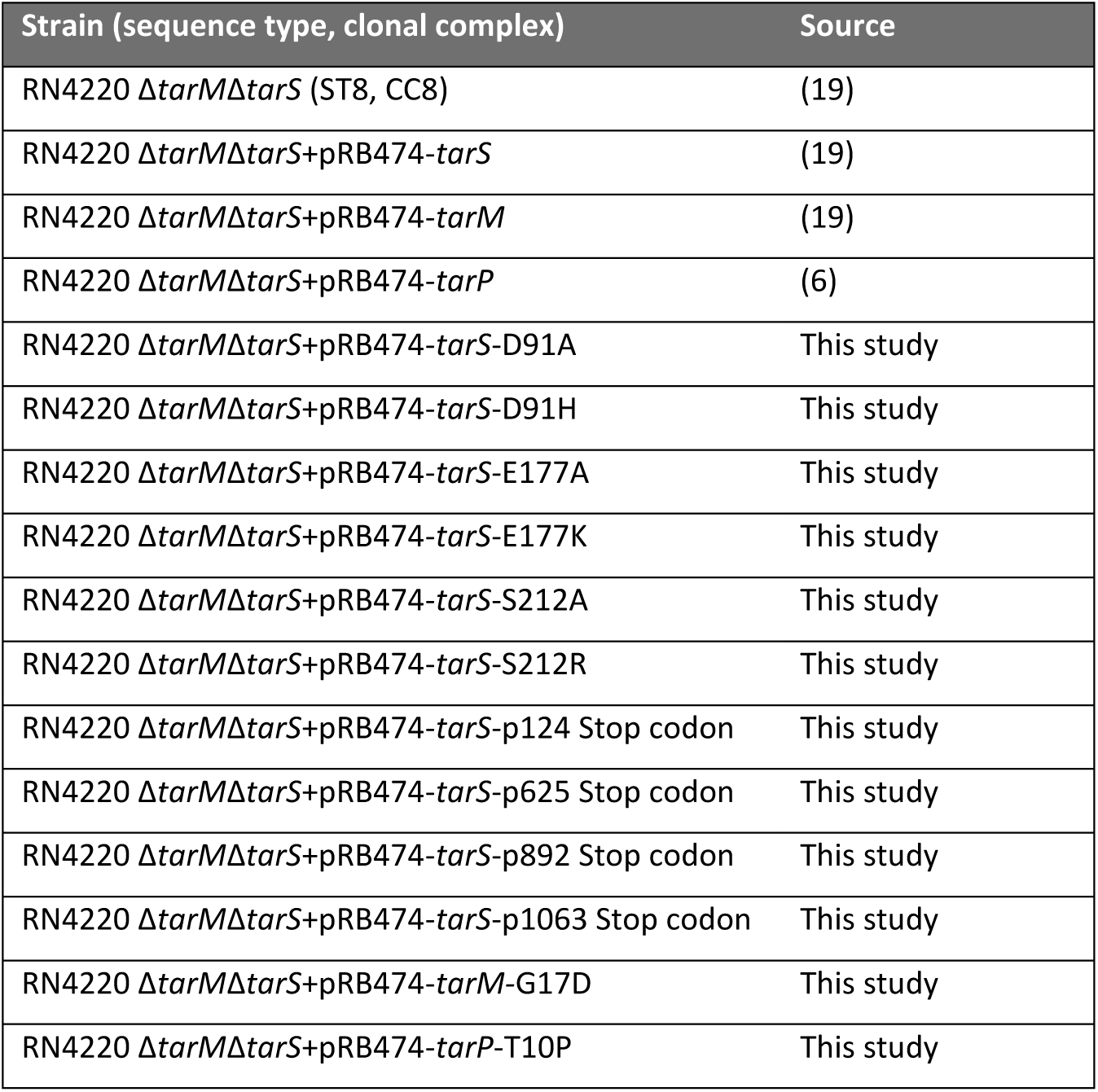
All bacterial strains used in this study

| Strain (sequence type, clonal complex) | Source |
| --- | --- |
| RN4220 $\Delta tarM\Delta tarS$ (ST8, CC8) | (19) |
| RN4220 $\Delta tarM\Delta tarS$+pRB474-*tarS* | (19) |
| RN4220 $\Delta tarM\Delta tarS$+pRB474-*tarM* | (19) |
| RN4220 $\Delta tarM\Delta tarS$+pRB474-*tarP* | (6) |
| RN4220 $\Delta tarM\Delta tarS$+pRB474-*tarS*-D91A | This study |
| RN4220 $\Delta tarM\Delta tarS$+pRB474-*tarS*-D91H | This study |
| RN4220 $\Delta tarM\Delta tarS$+pRB474-*tarS*-E177A | This study |
| RN4220 $\Delta tarM\Delta tarS$+pRB474-*tarS*-E177K | This study |
| RN4220 $\Delta tarM\Delta tarS$+pRB474-*tarS*-S212A | This study |
| RN4220 $\Delta tarM\Delta tarS$+pRB474-*tarS*-S212R | This study |
| RN4220 $\Delta tarM\Delta tarS$+pRB474-*tarS*-p124 Stop codon | This study |
| RN4220 $\Delta tarM\Delta tarS$+pRB474-*tarS*-p625 Stop codon | This study |
| RN4220 $\Delta tarM\Delta tarS$+pRB474-*tarS*-p892 Stop codon | This study |
| RN4220 $\Delta tarM\Delta tarS$+pRB474-*tarS*-p1063 Stop codon | This study |
| RN4220 $\Delta tarM\Delta tarS$+pRB474-*tarM*-G17D | This study |
| RN4220 $\Delta tarM\Delta tarS$+pRB474-*tarP*-T10P | This study |

**Supplementary Table 2.** Origin of WT pRB474 *tar*-gene inserts

| *Tar*-gene on plasmid RB474 | Genome of origin | Source | PubMLST allele |
| --- | --- | --- | --- |
| *tarS* | *S. aureus* RN4220 | (19) | 16 |
| *tarM* | *S. aureus* RN4220 | (19) | 1 |
| *tarP* | *S. aureus* N315 | (6) | 1 |

**Supplementary Table 3.**
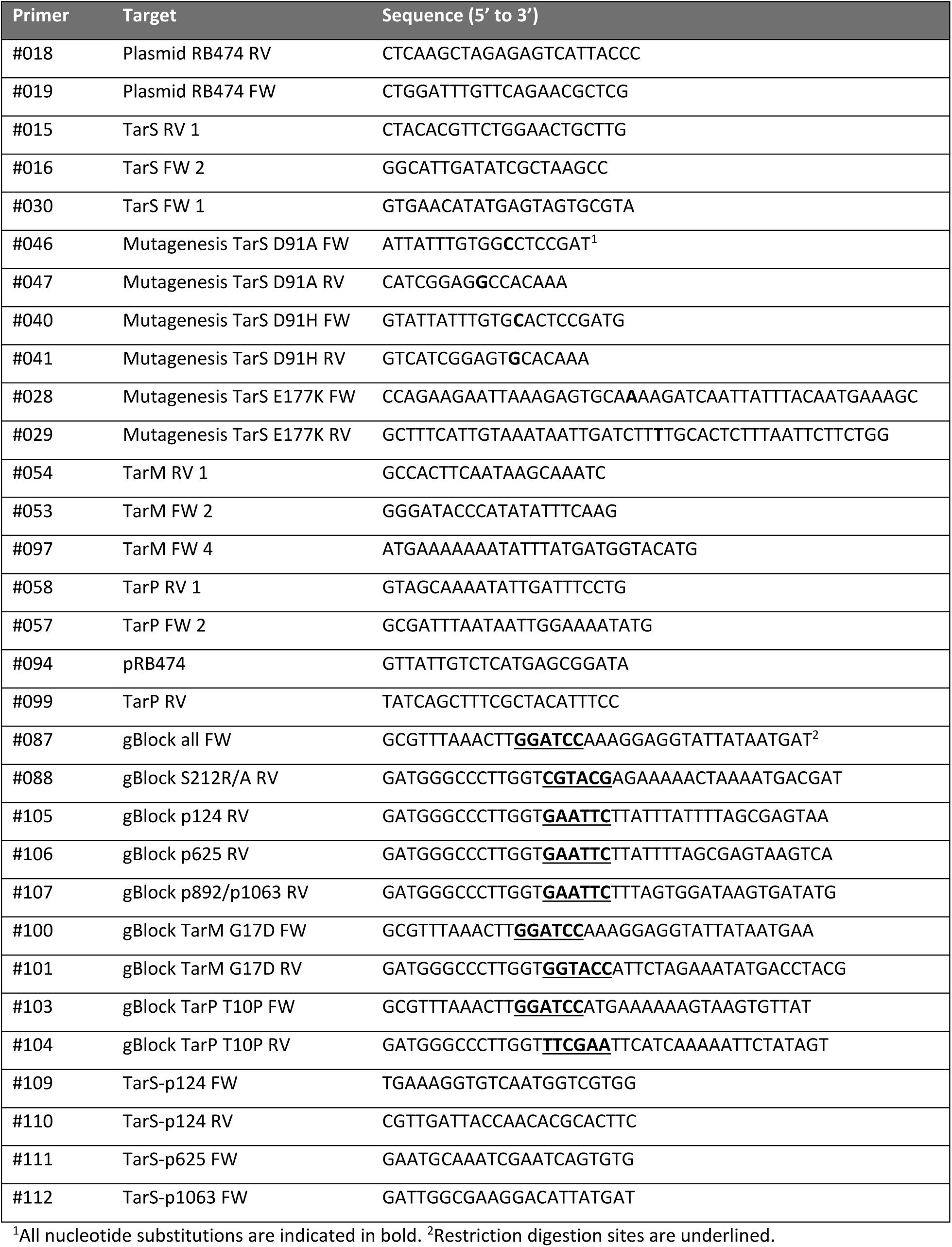
All primers used in this study

**Supplementary Table 4.** Number of *tar*-enzymes present in different clonal complexes

| CC | *tarS* | *tarM* | *tarP* |
| --- | --- | --- | --- |
| CC1 | 1,030 | 424 | 49 |
| CC5 | 5,761 |  | 1,205 |
| CC7 | 49 |  | 2 |
| CC8 | 6,661 | 6,600 | 15 |
| CC12 | 60 | 2 | 2 |
| CC15 | 436 | 398 | 3 |
| CC22 | 5,093 |  |  |
| CC25 | 96 |  |  |
| CC30 | 1,383 | 1,351 | 3 |
| CC45 | 584 |  | 38 |
| CC59 | 154 |  | 2 |
| CC88 | 111 |  | 5 |
| CC93 | 446 | 446 |  |
| CC97 | 300 |  | 12 |
| CC121 | 132 | 108 |  |
| CC130 | 465 |  |  |
| CC133 | 247 |  |  |
| CC398 | 1,347 |  | 397 |
| CC425 | 141 |  | 2 |
| Unknown | 970 | 162 | 56 |
| Total | 25,466 | 9,491 | 1,791 |

**Supplementary Figure 1.**
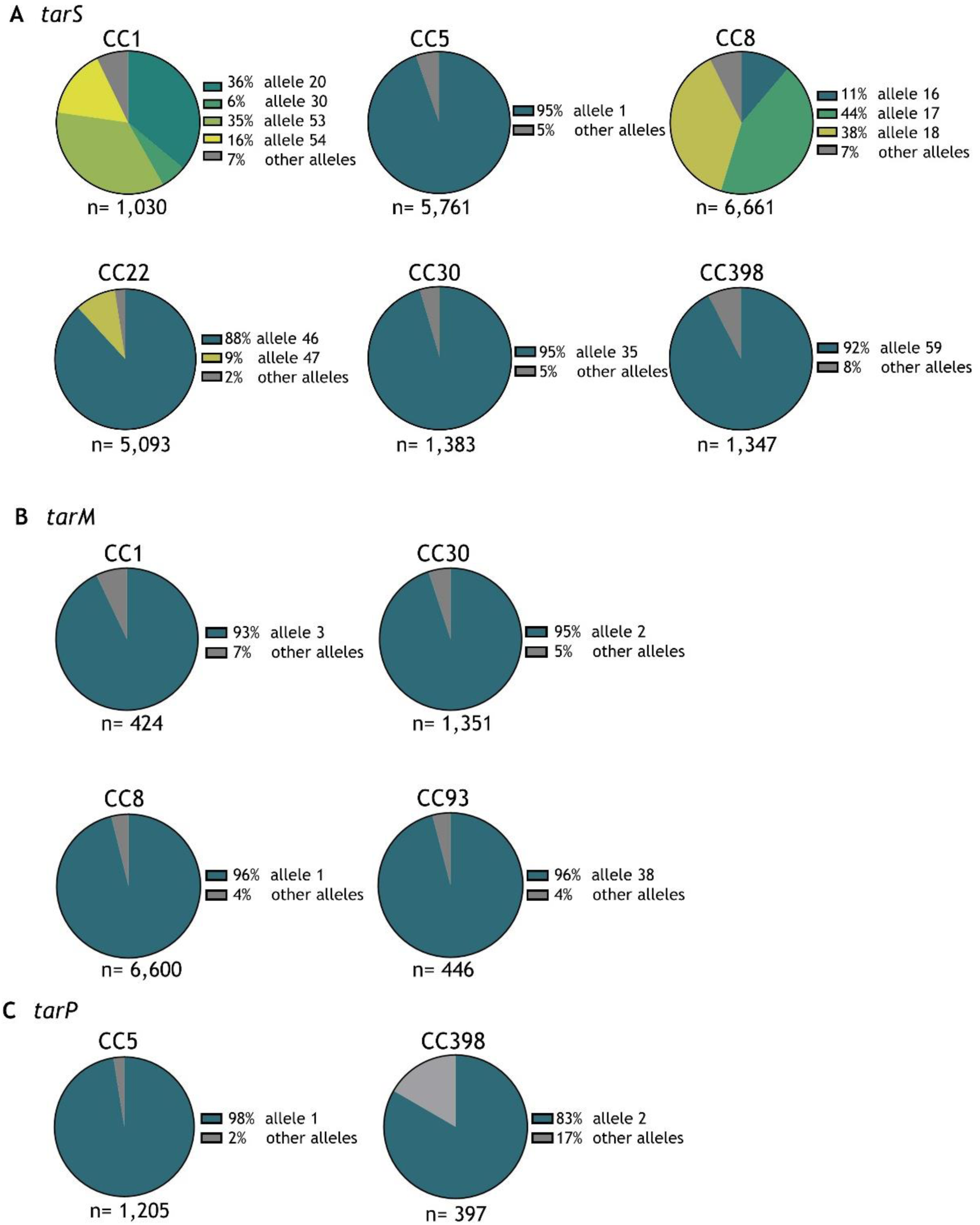
Distribution and occurrence of *tar* alleles within different clonal complexes. (A) The most frequently observed *tarS* alleles in distinct clonal complexes (CCs that comprise >4% of total isolates are depicted). (B) Same as in (A) but for *tarM*. (C) Same as in (A) but for *tarP*.

**Supplementary Figure 2.**
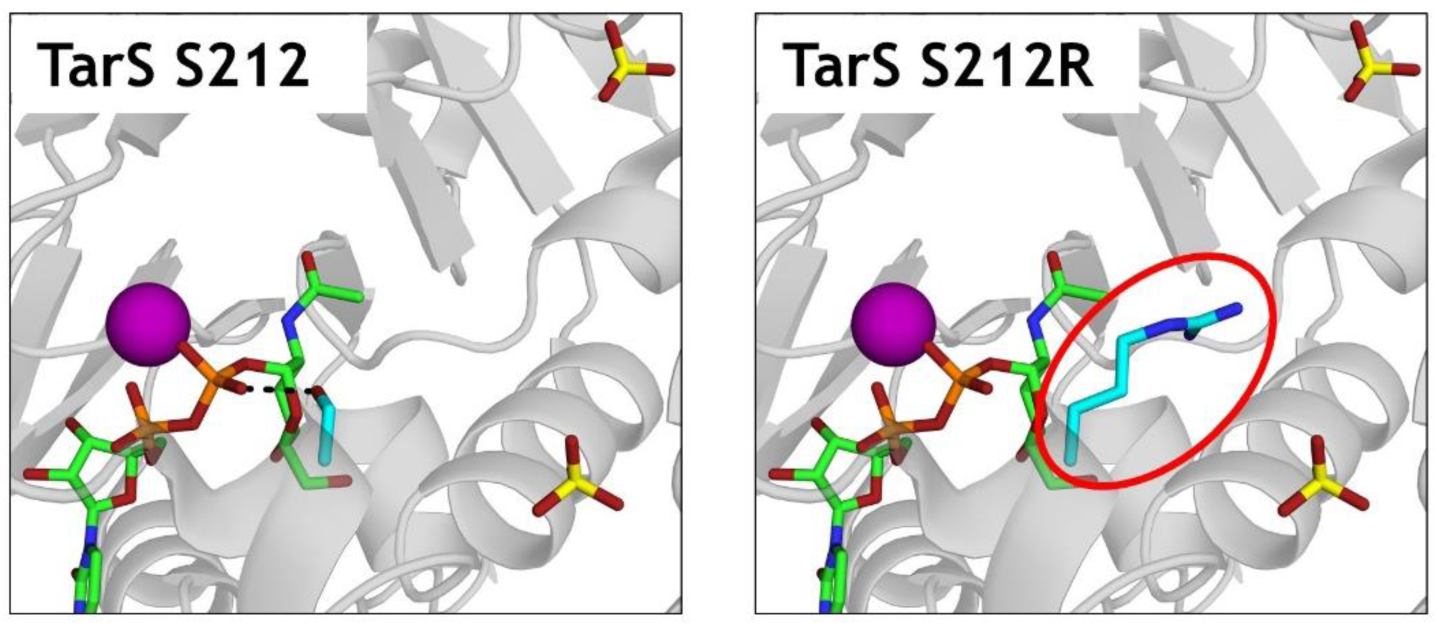
PyMOL visualizations of TarS S212 and S212R. TarS is shown in grey cartoon presentation (PDB code 5TZE). The donor substrate UDP-GlcNAc (green) and the residues of interest: S212 (left) and the substitution S212R (right, red circled), both in cyan, are all displayed in stick form. Sulfates are indicated in yellow.

